# The Role of Predictive Processing and Perceptual Load in Selective Visual Attention: An Examination with Semantically Salient and Less Salient Distractors

**DOI:** 10.1101/2024.10.15.618177

**Authors:** Ataol Burak Ozsu, Burcu A. Urgen

## Abstract

Our attentional resources are allocated to the various aspects of the environment based on the context, and predictive coding has been used as a model to explain the interaction between sensory-based information and top-down expectations in visual attention (Spratling, 2008; Rauss et al., 2011). On the other hand, the saliency of the environmental stimuli is also hypothesized to be capturing the attentional resources of the individuals involuntarily, and thus, it is thought to be playing a crucial role in attentional resource allocation. The current study investigates the role of predictive processing of task difficulty in selective visual attention in the presence of various distractors. Utilizing a letter search task, we provided brief cues about the upcoming task’s difficulty, and participants were asked to detect the target letters. We investigated whether predictive processing about task demands may cause a difference in behavioral measures in the presence of semantically less salient distractors in Experiment 1 (Gabor patches) and semantically more salient distractors in Experiment 2 (faces). Results showed that unmet expectations about the task demands caused longer reaction times in both studies. We observed that all independent variables, which are task difficulty, cue congruency, and distractor presence, affected reaction times in both experiments, but cue congruency interacted with distractor presence only in Experiment 2. Here, we argue that though predictive processing plays a role in attentional resource allocation and, distractors’ characteristics are also crucial as the saliency level interacts with the cue congruency.

## 1. Introduction

While processing a scene, the amount of information that can be captured by our senses is limited despite a wide array of environmental input, which exceeds the amount that can be processed by our brains at once. As a mechanism to navigate through this overwhelming environmental input, it is hypothesized that certain sensory input is selectively attended to while others are ignored, and this process is known as selective attention (Desimone & Duncan, 1995). Selective attention allows us to process the environmental input more efficiently by filtering the stimuli that are more relevant to the observer’s mental state. Based on the assumption that attention is selectively distributed depending on an individual’s goals and the contextual demands of the scene, one influential model is proposed, which is known as the “perceptual load theory” (Lavie, 1995). Studies conducted within this framework revealed that when the task demands are higher for the central task, meaning it requires significantly higher cognitive resources, the processing of task-irrelevant stimuli is hindered as fewer resources are available to process irrelevant stimuli, which in return causes less distraction (Lavie, 2005; Lavie, 2010). In contrast, when the perceptual load is low, the available cognitive capacity can be allocated to task-irrelevant stimuli, leading to higher distractions.

While the perceptual load theory has been widely utilized and accepted, there are also certain hypotheses that argue that the processing of task-irrelevant stimuli cannot be explained merely by the perceptual load required by the central task, but the characteristics of the distractors also play a significant role. One hypothesis that criticizes the perceptual load theory as it does not account for the identity of the distractors is the salience account proposed by Eltiti and colleagues (2005). It has been suggested that the perceptual salience of a distractor, or how attention draws a particular stimulus normally, could be a determinant of the distractor interference regardless of the perceptual load. Consequently, researchers found that when the salience of the distractors increases, their interference effect remains high even in the high perceptual load condition.

Building on the salience hypothesis, in the current study, we utilized distractors with varying semantic salience. In the original paper, salience was manipulated through temporal onset and the size of the distractor. While this approach evidently affected selective attention due to the alteration of low-level visual properties, how high-level visual properties affect distractor interference will be investigated as a part of the current study. Though several different distractors from diverse semantic categories have been used in perceptual load paradigms such as biological motion (Tunca et al., 2023), emotionally salient scenes (Hindi Attar & Müller, 2012; Gupta & Srinivasan, 2015), sounds (Murphy et al., 2013; Tellinghuisen & Nowak, 2003), and inanimate objects (Neumann et al., 2011), these studies used to test the effect of the perceptual load on the processing of the distractors not the effect of characteristics of the distractors on distractor interference, by comparing stimuli type that differs in their high-level properties. Consequently, in the current study, Gabor patches and faces are employed as distractors that differ in semantic salience. Gabor patches lack semantic content and would serve as a baseline for distraction that is primarily low-level rather than meaning-driven. In contrast, faces are regarded as visual stimuli that have great social and biological significance. Research suggests that inhibition of return can be used as a phenomenon to study if attention is reflexively allocated to a particular location since it only occurs under this particular condition, and faces are found to be the subjects of inhibition of return (Theeuwes & Van der Stigchel, 2006). By comparing these two types of stimuli, we aim to examine the effect of top-down influences on distractor interference. The current study aims to investigate the top-down processes of selective attention in the framework of perceptual load, and stimuli type composes one aspect of the current study that addresses the issue. Nevertheless, the perceptual load theory is a highly influential model of selective attention as it investigates selective attention through the manipulation of task demands. While perceptual load theory explains how task demands affect the allocation of cognitive resources, it does not fully account for the role of prior expectations in guiding attention. Predictive coding complements this by proposing that our cognitive system continuously generates expectations based on past experiences, shaping how we allocate attention before and after the stimuli are presented (de Lange et al., 2018). This study examines whether task demand expectations can facilitate selective attention in the presence of semantically salient and non-salient stimuli.

To be able to make sense of the sensory data gathered from the environment efficiently, one must not only process the input but also predict the most likely outcomes for quick inferences about the environment. As mentioned, the sensory input always contains lots of data to be processed, as well as ambiguities, which may cause inefficiencies in processing. Thus, certain templates to foster attention mediation are needed. As a model that would explain how our cognitive system handles the continuous excessive information flow, it has been proposed that we may utilize our prior experiences to efficiently process the sensory input (Spratling, 2017; Bar et al., 2006). Though this idea could be traced back to the works of Helmholtz (1867), who proposed that perceptual inferences are made according to the consistent features of the world, a more modern account has been provided by Rao and Ballard (1999) as predictive coding, which is a model that suggests visual processing is composed of feedback from higher cortical regions determining the activity in lower cortical regions whereas feedforward information from lower-level regions examine the match between predictions and the actual sensory information. Following research based on this theoretical framework has shown that expected stimuli are processed faster (Kok et al., 2012). Moreover, it has been argued that priors about the sensory input alter the perceptual processes (Sotiropoulos et al., 2011; Pinto et al., 2015). The predictive coding framework has been successfully utilized to address certain questions about perception by connecting prior knowledge and the actual sensory input.

Predictive processing has been employed to explain how attention is selectively modulated within a visual scene. This framework suggests that attention is guided by expectations regarding the location, timing, and features of stimuli, with attention being drawn to stimuli that match these expectations (Spratling, 2008). Similarly, modern accounts propose that neural activity following expected or unexpected stimuli is modulated based on whether these expectations are met (Walsh et al., 2020). Although the precise neural mechanisms underlying predictive processing are still under investigation, studies have shown that both neural activity and behavioral responses are influenced when feature-based and spatial attention is manipulated through expectation (den Ouden et al., 2010; Egner et al., 2010; Kok et al., 2012). However, these studies primarily focus on how predictions about specific stimulus features enhance decision-making accuracy and speed without fully considering how expectations about broader task demands and motivations might also shape attention. Indeed, a single theory cannot explain every aspect of visual perception. Nevertheless, the integration of complementary theories can add value to each other. Both perceptual load theory and predictive coding focus on how selective attention is guided, albeit from different perspectives—one driven by task demands and the other by expectations. Integrating these perspectives offers a more nuanced understanding of how attention operates.

A key distinction in predictive coding research is the difference between expectation and attention. While both can produce similar behavioral outcomes (Summerfield & Egner, 2009), they are conceptually distinct. Expectations involve predicting the likelihood of an event based on prior knowledge, whereas attention—specifically selective attention—focuses on the relevance of sensory input to a particular task or goal (Summerfield & de Lange, 2014). Therefore, when investigating how predictions influence attention, it is essential to clearly define and manipulate attention in the experiment and assess whether and how expectations impact attentional processes. Perceptual load theory is a useful tool for distinguishing these concepts as the attentional processes based on perceptual load manipulations are well-studied, and thus, the effects of expectations can be interpreted more reliably.

While perceptual load theory highlights how task demands drive attention during task execution, it overlooks how prior expectations, formed before the task and the expectation-based processes after the presentation of the task, shape this process. Predictive processing offers a complementary perspective by proposing that our cognitive system continuously generates predictions about sensory input based on prior knowledge, guiding selective attention accordingly. Integrating these two theories can be beneficial for a more comprehensive understanding of how top-down processing—particularly task demand expectations—affects selective visual attention, especially when the characteristics of task-irrelevant distractors are controlled.

Building on the integration of perceptual load theory and predictive coding, this study investigates how both task demands and prior expectations about those demands shape selective attention. Specifically, we examine the timing and accuracy of behavioral responses in two experiments designed to test the interaction between perceptual load, expectation, and the salience of distractors. The only difference between these experiments was the peripheral task-irrelevant distractors. The distractors in Experiment 1 were Gabor patches, which were utilized since the semantic information contained by Gabor patches is limited, whereas in Experiment 2, faces, which are hypothesized to be distractors with significantly more information carrying than Gabor patches, were presented as task-irrelevant distractors. While the paradigm itself clearly distinguishes the task relevance of the distractors, which allows us to study the effect of expectations on selective attention, we also aimed to investigate whether any effect observed in one experiment may persist when the saliency of the distractors is altered. We hypothesized that when the cues about the task demand prime a lower cognitive load for the central task, but the actual task demand is higher, the mean reaction time would increase while accuracy rates would decrease in comparison to congruent hard trials in which cues refer to a high cognitive load trial and the actual task demand is also high. However, when the cue indicates a harder task (high cognitive load) is about to be presented, but the actual task demand is lower, we expect to observe no difference between this condition and the condition in which the cue correctly predicts the lower task demand. Because the “hard cue” is hypothesized to cause an expectation that would mediate the selective attention toward the central task. Furthermore, we expect to observe an interaction effect between the cue congruency and the distractor presence in Experiment 2 (faces) on mean reaction times since the attention guided by expectations would cause an advantage when other stimuli that capture attention are presented and we hypothesized that lack of semantic saliency of Gabor patches would make them less attention capturing and the interaction effect between the cue congruency and distractor presence would not be observed in Experiment 1 (Gabor patches). While previous hypotheses are the hypotheses about the novelties of the current study, based on the previous literature, we also expect to observe the effect of cognitive load, distractor presence, and the interaction effect between cognitive load and distractor presence in both of the experiments.

## 2. Methods

### 2.1.1 Participants

#### Experiment 1

A total of 20 participants (16 female, Mean age = 21.8, age range [18 31]) with normal or corrected-to-normal vision participated in the study. The number of participants was determined based on the power analysis conducted with G*Power 3 (Faul et al., 2007) prior to the experiment (alpha = 0.05, 1-alpha = 0.95, effect size .3). None of the participants had a history of clinically diagnosed neurological or psychiatric disorder and were not using any psychiatric medication. The Human Research Ethics Board of Bilkent University approved the study. Participants were informed about the experimental procedure both verbally and through a written document, and they signed consent forms after they were given the information about the experiment. After the experiment, participants were compensated for their time with course points.

### 2.2.1 Stimuli, experimental design, and procedure

The visual search task (Lavie & Cox, 1997) was utilized to manipulate the perceptual load, which also allowed us to provide predictive cues about task demands on a trial-by-trial basis. Two kinds of stimuli were presented in the experiment. The distractors on the periphery were Gabor patches (1) created through a publicly available Gabor-patch generator code by Sebastiaan Mathôt (http://www.cogsci.nl/software/online-gabor-patch-generator). Four different orientations (45, 90, 135, and 180 degrees) were used to prevent any interactions between the letter location and the Gabor patch orientation. Other parameters contributing to the creation of Gabor patches held constant across different orientations: Standard deviation: 12 pixels, frequency in cycles/pixels: .1, phase: 0. The visual degree values for every Gabor patch from the center of the screen was 13 degrees. The letters and the fixation point (2) presented in the center of the screen were generated and presented via Psychtoolbox (Kleiner et al., 2007) on MATLAB /The Mathworks, Natick, MA).

The visual search task consisted of consecutive presentations of cues about the upcoming task’s difficulty, the task itself, which required the participants to detect the target letters X and N among five other letters, and feedback about their accuracy on the task. The task was adapted from the original study when the procedure was initially presented (Lavie & Cox, 1997), and the following studies used this paradigm to manipulate perceptual load and the other features of the paradigm, such as distractors’ distance to the central task, (Theeuwes et al., 2004) priming effects such as spatial cues about the target letter location (Johnson et al., 2002). Here, by utilizing the flexibility provided by the procedure, we present cues, task-irrelevant distractors, and feedback about the accuracy in addition to the original version of the task. The cue was presented to participants through the experiment screen, which was either HARD which is a cue signing that the upcoming central task would be a high perceptual load task or EASY which is a cue signing that the upcoming central task would be a low perceptual load task, and the size and the font (Helvetica) of the letters were identical to the letter sizes presented in the visual search task. The cues were presented for a duration of two seconds, which was determined based on the pilot studies conducted. In 75% of the trials, the cue correctly predicted the perceptual load. The letters were presented for one second, which was also based on the pilot studies and the pilot participants’ average response times, and the feedback phase was also presented for 1 second. In the low-load condition, the letters except the target letter were Os, whereas in the high-load condition, the letters except the target letter were randomly selected from the [H, W, V, K, M, Z] pool. In both conditions, participants were instructed to press X if they detected X and press N if they detected N on the screen as accurately and as fast as possible. The task of the participants, the number of letters presented, and the durations did not differ across trials and blocks. The task difficulty manipulation was randomly distributed to prevent any expectation biases while the target letter appearance, the peripheral side where distractors were presented, and the orientation of the Gabor patches were counterbalanced. Each Gabor patch was presented 96 times, which constituted half of the trials, as in 384 trials, no distractors were presented. Overall, the experiment consisted of 768 trials, which were divided into eight blocks.

A practice task was provided to each participant to familiarize them with the task. All participants were instructed to fixate on the center -the position where the fixation point was presented- and they were given the information that the cues only correctly predict the perceptual load in 75% of the trials. The screen was viewed at a 56 cm distance.

#### Experiment 2

### 2.1.2 Participants

A total of 20 participants (17 female, Mean age = 22.1, age range [18 43]) with normal or corrected-to-normal vision participated in the study. The number of participants was determined based on the power analysis conducted with G*Power 3 (Faul et al., 2007) prior to the experiment (alpha = 0.05, 1-alpha = 0.95, effect size .3). None of the participants had a history of clinically diagnosed neurological or psychiatric disorder and were not using any psychiatric medication. The Human Research Ethics Board of Bilkent University approved the study. Participants were informed about the experimental procedure both verbally and through a written document, and they signed consent forms after they were given the information about the experiment. After the experiment, participants were compensated for their time with course points.

### 2.2.2 Stimuli, experimental design, and procedure

Experiment 2 was identical to Experiment 1 in terms of cue, task, and feedback durations, task demands, and other parameters except for distractors. In this experiment, four male and four female faces were used as distractors. The faces were taken from the Chicago Face Database (Ma et al., 2015). Only white models were used to prevent any semantical difference between models, which may affect the distraction power. Only white models were presented since if models from different races were presented, it could have caused variability between distractors on a level that falls outside of the scope of this experiment. Also, only faces with neutral facial expressions were utilized to ensure that facial expressions may not cause another semantic dimension to be taken into account. Though it was attempted to control for the overall size within faces and between Gabor patches from Experiment 1, the shape of the faces caused negligible size differences. The visual degrees for faces were identical to Gabor patches (13 degrees from the center) but the shape differences between faces and the circle shape of Gabor patches caused slight differences in terms of the area occupied within their positions. Also, we controlled for the low-level features of distractors between studies and within this study by using the SHINE toolbox (Willenbockel et al., 2010). Initially, the background/foreground segmentation was done through a custom Python code both for Gabor patches and the faces, then we performed luminance correction with the SHINE toolbox. We did not perform the contrast correction as it caused a significant distortion to faces and their overall shape.

#### 2.3 Data collection and analysis

The response times and accuracy data for all congruency, perceptual load, and distractor presence were collected and analyzed for both studies. Response time was defined as the time interval between the onset of letters on the screen and the time point that the participant pressed either X or N. If the participant failed to give a response in a 1 second period, the trial was marked as a missed trial. The accuracy rate indicates the portion of the trials in which the participant correctly pressed the target letter button that appeared on that particular trial. Only the trials that were not missed were analyzed for accuracy and response time. For both experiments, we conducted repeated measures analysis of variance (ANOVA) to investigate how cue congruency (congruent, incongruent), distractor presence (present, absent), and perceptual load (high, low) affected the mean response time and accuracy.

## 3. Results

### 3.1 Experiment 1

The analysis of reaction times illustrated that participants took longer to respond when the cue about the perceptual load did not match the actual perceptual load. Repeated measures ANOVA (2 (Perceptual load) x 2 (Congruency) x 2 (Distractor availability)) demonstrated a main effect of congruency on the mean response time (F [1,19] = 7.592, p = 0.013, ηp2 = 0.285) (See Fig.2.a). However, this effect did not persist in the accuracy analysis (F [1,19]= 4.741* 10^-4^, p= 0.983)

**Figure 1.**
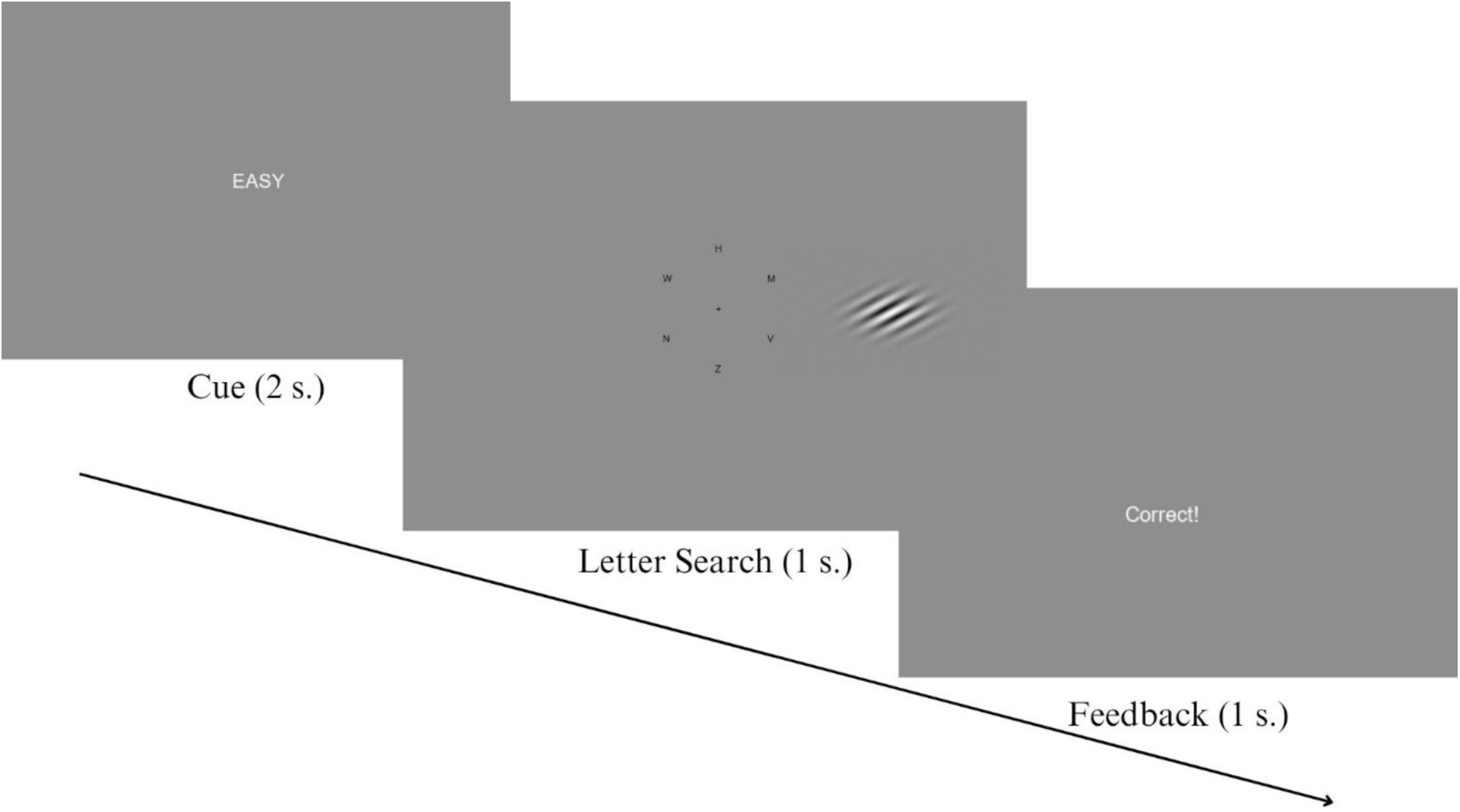
The experimental flow illustrating an incongruent trial with high perceptual load and a correct response in Experiment 1, in Experiment 2, instead of Gabor patches, faces were used as distractors.

**Figure 2.**
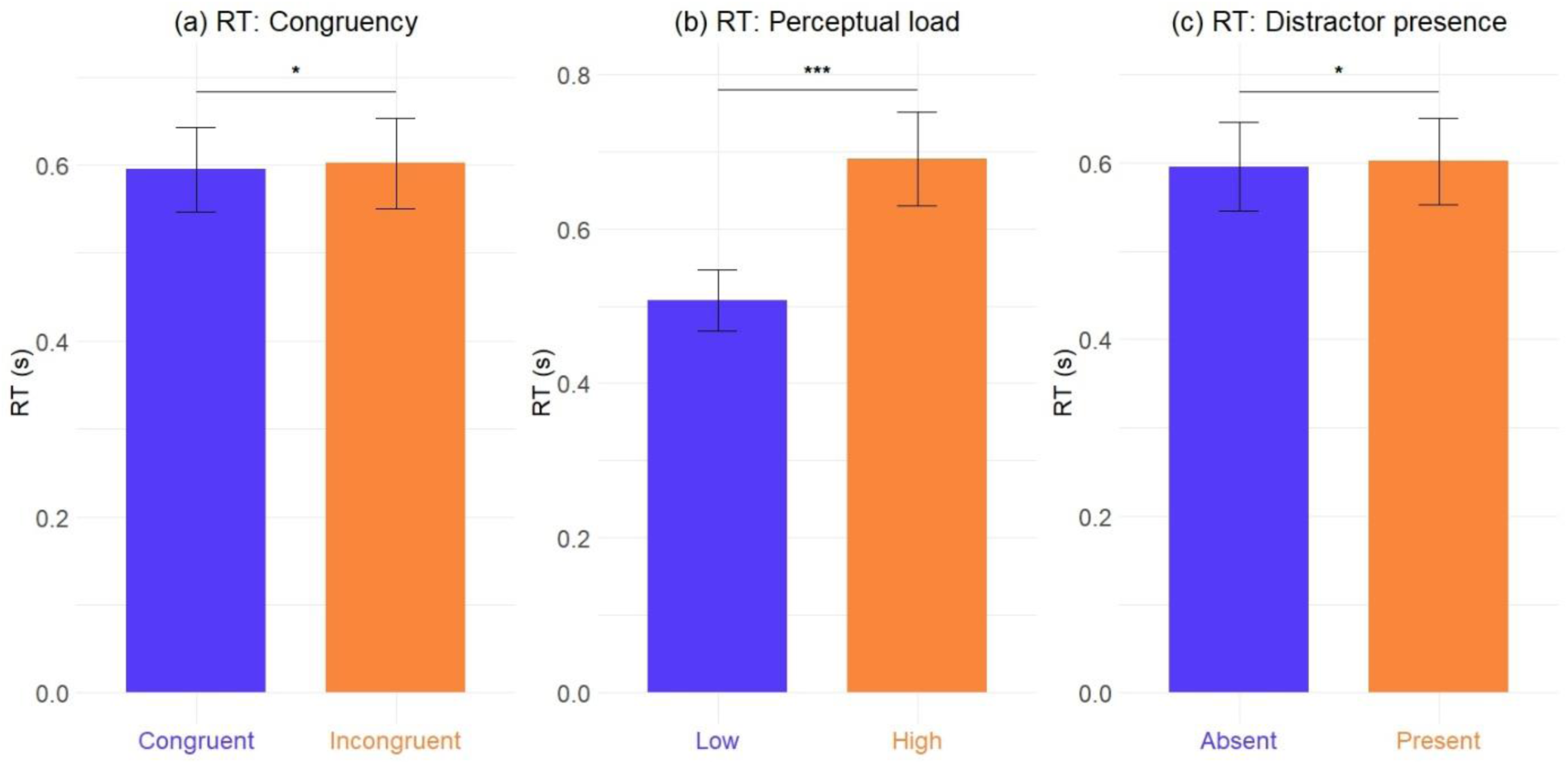
The mean reaction time comparison for congruency (a), perceptual load (b), and distractor presence (c) for Experiment 1.

Additionally, participants took longer to respond when the perceptual load was higher. The repeated measures ANOVA revealed the main effect of the perceptual load on mean response time (F [1,19] = 263.319, p < 0.001, ηp2 = 0.933) (See Fig. 2.b.). Additionally, the effect of the perceptual load was also observed in the accuracy analysis (F [1,19] = 51.740, p < 0.001, ηp2 = 0.731)

Distractors’ presence followed a similar trend to the cue congruency. Participants responded more slowly in the presence of Gabor patches compared to the trials where distractors were absent (F [1,19] = 8.409, p = 0.013, ηp2 = 0.285) (See Fig. 2.c.). However, the presence of Gabor patches did not affect the accuracy rates of the participants (F [1,19] = 8.433* 10^-4^, p = 0.977)

In terms of mean reaction time, other than the distractor presence and congruency interaction, our analyses did not reveal any significant interaction effects. However, we observed a marginal interaction effect between distractor presence and congruency (F [2,38] = 3.173, p = 0.081, ηp2 = 0.152). Perceptual load and distractor presence (F [2,38] = 1.422, p = 0.248), perceptual load and congruency (F [2,38] = 2.074, p = 0.166), and the three-way interaction between the independent variables perceptual load, distractor presence and congruency (F [3,57] = 0.441, p = 0.515) were insignificant.

In terms of accuracy rate, all interaction effects were insignificant, as the statistics were calculated as follows: Perceptual load and congruency interaction (F [2,38] = 1.601, p = 0.221), perceptual load and distractor presence interaction (F [2,38] = 0.030, p = 0.864), congruency and distractor presence interaction (F [2,38] = 0.265, p = 0.613) and the three-way perceptual load, congruency, distractor presence interaction (F [3,57] = 0.126, p = 0.727) were insignificant.

### 3.2 Experiment 2

An identical repeated measures ANOVA (2 (Perceptual load) x 2 (Congruency) x 3 (Distractor presence)) to Experiment 1 demonstrated a similar pattern for both reaction times and accuracy rates in Experiment 2. Participants responded significantly slower in the incongruent trials in comparison to congruent trials (F [1,19] = 30.063, p = 0.002, ηp2 = 0.613) (See Fig. 3.a). However, the congruency variable did not affect the participants’ accuracy in general (F [1,19] = 1.257, p = 0.276)

**Figure 3.**
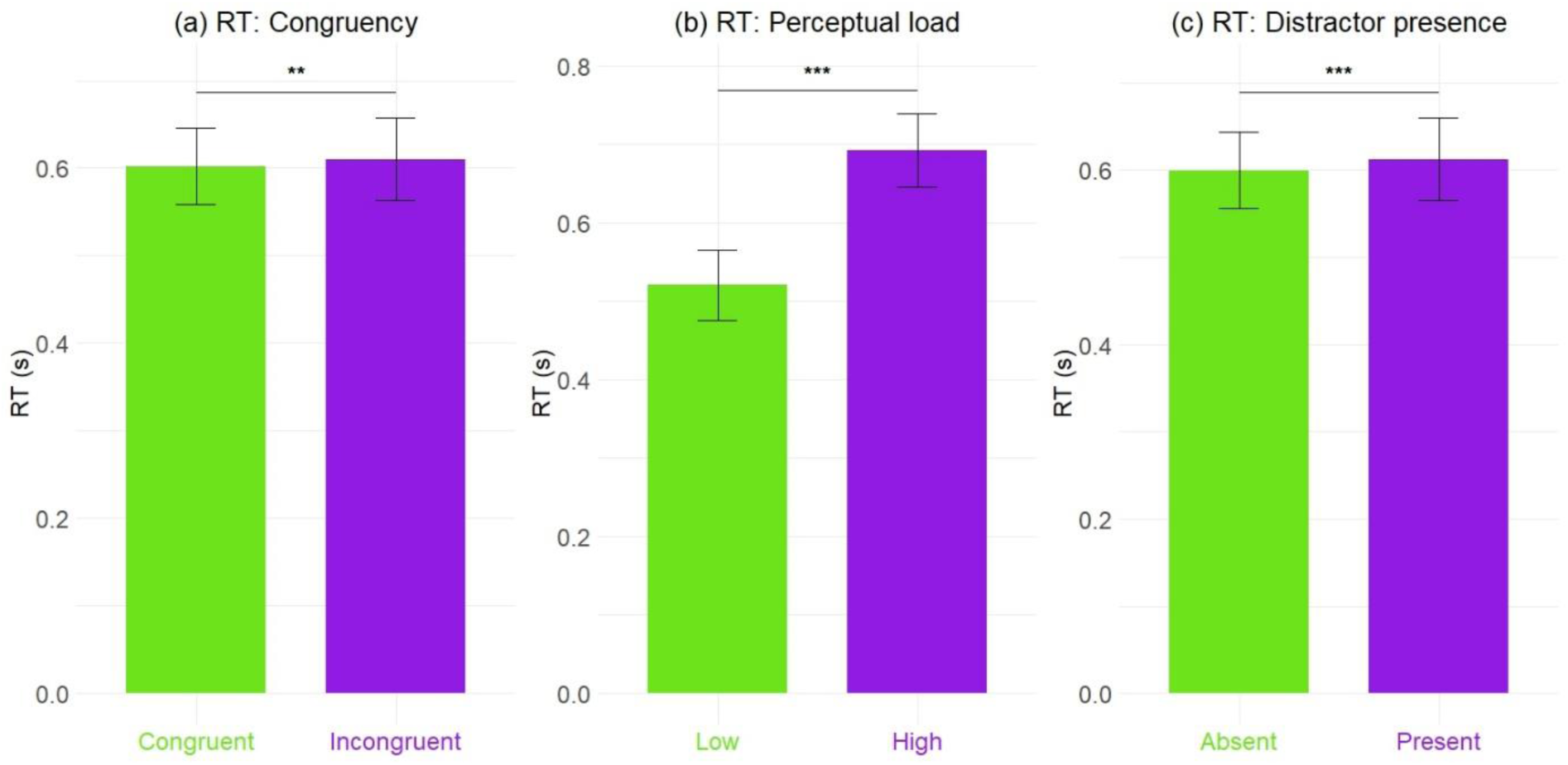
The mean reaction time comparison for congruency (a), perceptual load (b), and distractor presence (c) for Experiment 2.

In Experiment 2, in the low perceptual load condition, participants gave significantly faster responses compared to the high perceptual load condition (F [1,19] = 686.240, p < 0.001, ηp2 = 0.972) (See Fig. 3.b). Furthermore, the accuracy rates were significantly lower in the high perceptual load trials (F [1,19] = 105.996, p < 0.001, ηp2 = 0.653).

In Experiment 2, the effect of the distractors resulted in a similar effect to Experiment 1. The presence of distractors caused delayed responses in comparison to trials with distractor absent display condition (F [1,19] = 30.695, p < 0.001, ηp2 = 0.618) (See Fig. 3.c). On the other hand, the accuracy rates did not differ between distractor present and absent conditions (F [1,19] = 0.299, p = 0.693).

In addition to the main effects, our analysis revealed an interaction between the variables distractor presence and congruency (F [2,38] = 9.392, p = 0.006, ηp2 = 0.331) (See Fig. 4). Subsequent post hoc analyses demonstrated that while incongruent cues resulted in delayed responses after Bonferroni correction in the presence of faces (t=4.758, p_bonf_< 0.001) when no distractor was presented, the cue congruency did not cause any significant changes in terms of mean reaction times (t=0.346, p_bonf_=1.000).

**Figure 4.**
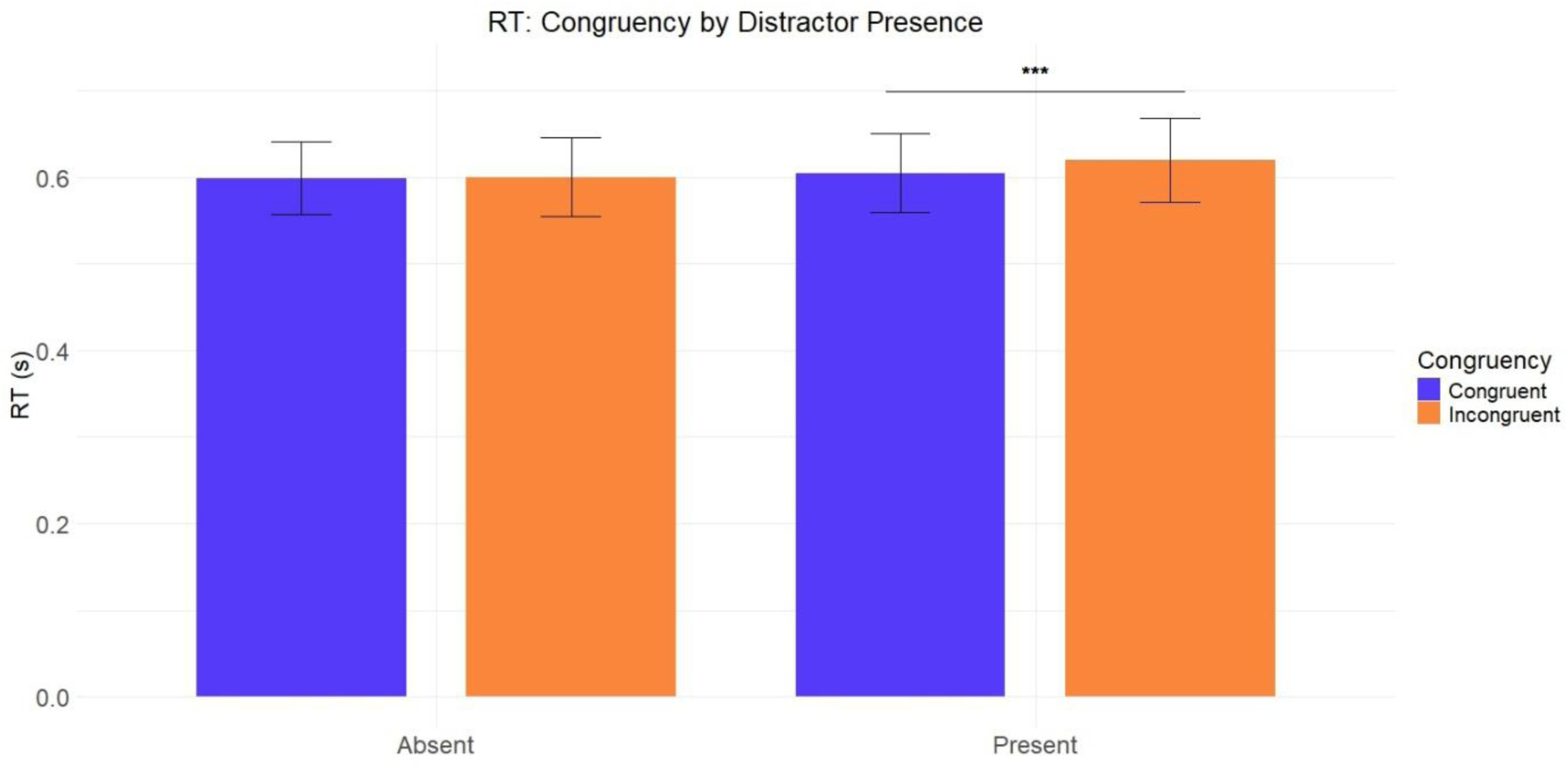
The mean reaction time comparison for the congruent and incongruent trials when grouped for distractor presence.

In terms of mean reaction time, other than the distractor presence and congruency interaction, our analyses did not reveal any significant interaction effects. However, we observed a marginal interaction effect between perceptual load and congruency (F [2,38] = 3.173, p = 0.091, ηp2 = 0.143). Perceptual load and distractor presence (F [2,38] = 0.009, p = 0.924) and the three-way interaction between the independent variables perceptual load, distractor presence, and congruency (F [3,57] = 0.886, p = 0.358) were insignificant.

In terms of accuracy rate, all interaction effects were insignificant, as the statistics were calculated as follows: Perceptual load and congruency interaction (F [2,38] = 0.333, p = 0.571), perceptual load and distractor presence interaction (F [2,38] = 0.541, p = 0.471), congruency and distractor presence interaction (F [2,38] = 0.071, p = 0.793) and the three-way perceptual load, congruency, distractor presence interaction (F [3,57] = 0.017, p = 0.897) were insignificant.

### 3.3 Between Experiment Comparison

An independent samples t-test was conducted to compare the effects of congruency, perceptual load, and distractor presence between studies.

#### 3.3.1 Congruency

We compared the time differences between incongruent and congruent trials between studies through an independent samples t-test. In Experiment 2, the effect of the cue congruency was significantly higher than in Experiment 1 (T (38) = 3.435, p = 0.001, Cohen’s d= 1.086) (See Fig 5.a).

**Figure 5.**
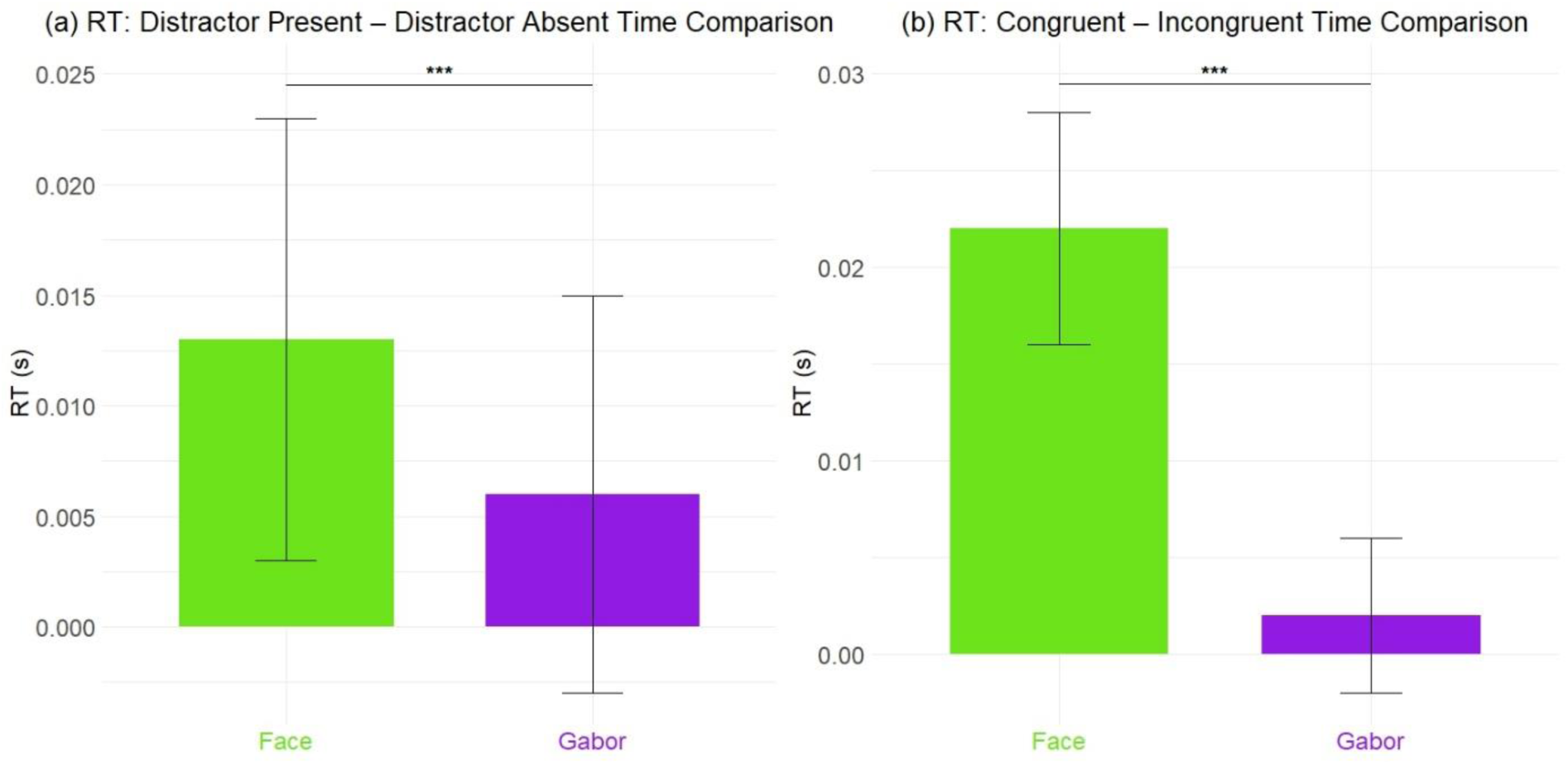
The mean reaction time difference comparison between experiments for distractor presence (a), congruency (b)

#### 3.3.2 Perceptual Load

We compared the time differences between high perceptual load and low perceptual load trials between studies through an independent samples t-test. We conducted Welch’s t-test as the assumption of equal variances was violated as Levene’s test yielded significant results. There were no significant differences between studies for the reaction time differences between high-load and low-load studies. (T (38) = 0.798, p = 0.431, Cohen’s d= 0.319).

#### 3.3.3 Distractor presence

We compared the time differences between distractor present and distractor absent trials between studies through an independent samples t-test. In Experiment 2, the effect of the distractor presence was significantly higher than in Experiment 1 (T (38) = 2.171, p = 0.036, Cohen’s d= 0.334) (See Fig 5.b).

## 4. Discussion

The processing of sensory input relies on expectations and whether these expectations are matched. It is hypothesized that our brain utilizes prior information to make efficient perceptual inferences through several mechanisms, one of which could be moderating attention. On the other hand, attentional resource allocation has been a subject of discussion for a long period, and one long-lasting theory about it is the perceptual load theory. The current study investigates the effect of expectations regarding the perceptual load on behavioral responses as the perceptual load theory provides a framework that allows us to investigate how priors about a particular task affect the processing of the visual input. The perceptual load paradigm clearly distinguishes the task and the task-irrelevant features, which could provide information about how our attention is modulated between those features. Hence, we conducted two studies that only differ in terms of the distractors’ semantic saliency, which allowed us to investigate how the distractors’ characteristics affected the attentional resource allocation based on expectations, which is overlooked in perceptual load studies.

Depending on the predictive processing framework, we hypothesized that when the expectations about the perceptual load of a given task are not matched, the reaction times would increase as the pre-stimulus onset information is not useful to process the input. Nevertheless, when the cue indicates a high-load task, but the task is actually low-load, we hypothesized that, in both studies, the processing should not be hindered as much as it would in a trial in which the cue indicates a low-load task, but the task is actually high-load. When the cue indicates a high-load task, it is hypothesized that participants would direct their attention to the central task more in comparison to the low-load cue, and as a result, the reaction times and accuracy rates should not be impaired when the high-load cue is presented. Moreover, we posited that as the faces possess more semantic knowledge due to their nature, when the expectations about the task are not matched in Experiment 2, we would expect a larger temporal delay to respond compared to Experiment 1 as the semantically more salient stimuli are hypothesized to compete more for the attention of the observer. In addition to the aforementioned study-specific hypotheses, we expected to observe the effects of perceptual load, distractor presence, and the interaction between perceptual load and distractor presence in both of the studies, as both effects were replicated several times in the existing literature.

### 4.1 How prior information about perceptual load mediates attention

Though there is a convincing amount of prior work that suggests that unmet expectations cause a significant decrease in task performances, most of the studies utilize spatial (Aitken et al., 2020), temporal (Vangkilde et al., 2012), or feature-based cues (Stein & Peelen, 2015) about the task. Here, as a novelty, we take advantage of the clear distinction between task and task-irrelevant features in the perceptual load paradigm, and through building expectations about the task demands, we investigate attention modulation through expectations. In both of the experiments, when the predictions about the upcoming task demands did not match the actual task demands, participants took longer to respond. One explanation could be that the priors allowed participants to better allocate their attentional resources before stimulus onset, resulting in reduced reaction times. Another explanation might be that prediction errors regarding the priors presented to the participants resulted in poor attention mediation, which causes a re-allocation process of resources after the stimulus onset that may be time costly. Furthermore, both of these hypotheses may not be mutually exclusive, and both of them might have affected the reaction times of the participants. The underlying processes could be better studied with other tools and paradigms. In the current study, the priors we presented had a 75% chance of being a correct indicator of the following task’s perceptual load. The predictive processing algorithms rely on the computations that determine the weight of the prior information as determinants of the predictions based on their validity (Spratling, 2017). Consequently, when the larger portion of the priors correctly predicts the actual sensory input, the reliance on the prior information increases. As a result of that, in the current study, the participants are relying on the priors more in comparison to a hypothetical %50 validity experiment. Therefore, the weight of expectations on perceptual inferences in the current study is larger than that of conventional perceptual load experiments. The results show that expectations about the task demands affect the perceptual inferences, which are compatible with the framework, but we observed small differences, though significant. This can be attributed to a learning process in which participants realize relying on the cues may not necessarily enhance their reaction times. However, our analyses revealed another important aspect that amplified the differences between incongruent and congruent trials, which is the semantic saliency of the distractors.

### 4.2 Distractor’s semantic saliency and its interaction with predictive processing

The perceptual load paradigms have been criticized for the issue that the distractor interference level is mediated by the saliency of the distractors, referring to the baseline level of attention attraction of the distractors (Eltiti et al., 2005). In the current study, as we aim to investigate how visual attention is mediated by expectations, we aim to control this problem. We created two experiments that only differed on the semantic saliency of the distractors and controlled for the low-level features. Initially, in Experiment 2, we observed that the cue incongruencies resulted in higher reaction times when the faces were presented compared to no-distractor trials when the cue did not correctly predict the actual load, which gives a clue about how our cues were utilized. Expectation mismatches may affect behavioral responses more when there are features on the visual field that compete for attention capture because focusing an adequate amount of attention on the central task may be more difficult when the distractors compete more for attention. To further explore this effect, we compared the effect of cue congruency on response times in between experiments. The results illustrated that the reaction time difference between congruent and incongruent trials in Experiment 2 is significantly larger compared to Experiment 1, which supports the view that the unmet expectations cause higher delays when the task-irrelevant features are more attention-capturing. The findings of this study demonstrate that predictive processing is involved in perceptual load-based attention mediation, and this effect interacts with the characteristics – semantics in this case-of these features that compete for the attentional resources of an individual.

## 5. Conclusion

Our investigation of the effects of priors about the perceptual load on the processing of the task and the task-irrelevant distractors showed that the visual attention mediation through expectations about the perceptual load required by the central task interacts with the semantic saliency of the distractors. Though it is well-studied that predictions about particular scenes are prominent in how these scenes are processed, the current study examines the effect of the semantics of the task-irrelevant features and utilizes the perceptual load paradigms to study the interaction between expectations and selective attention, which can contribute to our general knowledge about predictive processing in mind. Though these findings provide certain insights about the interplay between predictions and attention, they also illustrate the need for studies that may reveal more about the underlying mechanisms of this interaction. Although widely utilized, the perceptual load paradigm utilized in this study lacks ecological validity. Furthermore, the current study provides very limited information about the underlying processes of the expectations of task demands. Neuro-imaging studies, naturalistic scenes, and eye-tracking methods can all be utilized to get a more comprehensive understanding of how predictive processing affects attention in various contexts and illuminate what happens from the beginning point of building predictions to the point where perceptual inferences are made.

